# Transcriptional Reprogramming Drives Cold Adaptation During Long-Term Starvation in *Saccharomyces eubayanus*

**DOI:** 10.1101/2025.11.19.689257

**Authors:** Luis A. Saona, Tamara Mateluna-Cáceres, Pablo Villarreal, Macarena Las Heras, Vasni Zavaleta, Francisco A. Cubillos

**Affiliations:** Universidad de Santiago de Chile, Facultad de Química y Biología, Departamento de Biología, Santiago, Chile; Millennium Nucleus of Patagonian Limit of Life (LiLi), Valdivia, Chile; Centro Científico y Tecnológico de Excelencia Ciencia & Vida, Fundación Ciencia & Vida, Huechuraba, Santiago, Chile; Millennium Institute for Integrative Biology (iBio), Santiago, Chile; Centro de Genómica y Bioinformática, Facultad de Ciencias, Ingeniería y Tecnología, Universidad Mayor, Santiago, Chile

**Keywords:** Aging, Long-term cultures, Low-temperatures, *S*. *eubayanus*

## Abstract

The ability of microorganisms to survive prolonged periods of nutrient scarcity is essential for their survival. Yet, the underlying adaptive mechanisms remain partly understood, especially in non-model eukaryotes. Here, we examined how the cryotolerant yeast *Saccharomyces eubayanus* adapts to 60 days of cold (4°C) starvation, focusing on the roles of genetic and transcriptional changes. We find that the primary engine of the long-term adaptation is a stable, reprogrammed transcriptional state, rather than the selection of point mutations. Aged isolates exhibited improved growth performance and cryotolerance, a phenotype that remained stable for ∼40 generations. This specialist adaptation involved a trade-off with tolerance to other stresses. Whole-genome sequencing revealed few fixed mutations, indicating that genetic variation did not drive the phenotype. Transcriptomic analysis revealed a significant physiological reprogramming, with cells shifting from anabolic activity to a catabolic, scavenging state driven by enhanced respiration and activation of the General Stress Response. This work highlights that a stable transcriptional state drives long-term cold adaptation, providing the foundation for the superior phenotype of aged isolates that persist through generations.

## Introduction

The ability to survive prolonged periods of stress and nutrient scarcity is a critical determinant of microbial fitness [1, 2]. In nature, microbial populations face low-nutrient cycles, imposing strong selective pressures that favour entry into dormant or quiescent states [1, 3]. During these phases of stillness, which can last for months or even years, cells must not only withstand the lack of nutrients but also other persistent environmental stresses, such as low temperatures [2, 4]. This scenario raises a fundamental evolutionary question: are these periods of dormancy merely passive, or do they represent an active period of selection and adaptation? In the bacterial world, the “Growth Advantage in Stationary Phase” (GASP) phenotype offers a plausible survival alternative [2, 5–10]. This phenomenon describes how mutants arise in aged cultures, which can grow in cell culture by utilizing nutrients released from their dead counterparts, eventually taking over the population [8]. This process of population “rejuvenation” is driven by the selection of mutations that optimise scavenger metabolism and stress resistance.

In eukaryotes, such as the yeast *Saccharomyces cerevisiae*, the study of ageing and long-term survival is addressed through the paradigm of Chronological Lifespan (CLS), defined as the time a non-dividing cell remains viable in stationary phase [11, 12]. CLS is an ecologically relevant framework for understanding how yeasts survive in the environment, and represents a model for ageing in higher organisms. Indeed, within these ageing populations, processes analogous to GASP can occur, where transcriptional plasticity and the selection of beneficial mutations drive the emergence of fitter individuals [13, 14]. Diploid *Saccharomyces* cells often respond to nutrient starvation by sporulating [15–17]. However, several studies show that not all cells or species follow this path. In natural environments where nutrient depletion and seasonal changes are common, yeasts often enter a quiescent state, which is a reversible, non-dividing phase marked by high stress resistance and reduced metabolism [18–22]. Evidence from wild isolates of *S. cerevisiae* shows that quiescent cells can persist for extended periods in soil, bark, or decaying fruit, resuming growth upon favourable conditions [23]. This suggests that quiescence, although less studied than sporulation, constitutes an ecologically relevant and evolutionarily stable survival strategy in the wild.

*Saccharomyces eubayanus* is the most cryotolerant species in the *Saccharomyces* genus and the progenitor of the industrial lager-brewing yeasts [24–32]. This yeast species represents an interesting system for studying adaptation to prolonged stress. Its natural habitat is the Patagonian forests, often linked with *Nothofagus* trees along the Andes treeline. These areas are marked by long and cold periods, indicating that these species likely developed effective strategies to survive under conditions of low temperature and nutrient scarcity [4, 33–35]. Previous studies in *S. cerevisiae* have shown that CLS long-term survival is an active process, driven by extensive transcriptional reprogramming towards a stress-resistant, quiescent state [12, 36, 37], often mediated by TORC1/PKA signalling [13, 18], and by the selection of advantageous mutations that allow fitter individuals to repopulate the culture [14, 38]. However, these key insights originate from experiments conducted at standard laboratory temperatures and conditions, which may not reflect the selective pressures of the natural cold environments inhabited by *S. eubayanus*. Therefore, critical gaps remain in our understanding: (i) how these adaptive mechanisms are deployed or potentially modified in a wild, cryotolerant species under constant stress of low temperature, and (ii) the relative contribution of genetic mutations versus stable transcriptional reprogramming to long-term microbial adaptation in such environments.

To address this question, we used a 60-day chronological lifespan experiment at low temperature to investigate the adaptive dynamics of *S. eubayanus*. We hypothesized that long-term cold exposure would drive the emergence of fitter isolates and that this adaptation could be underpinned by both the selection of advantageous mutations and/or a stable reprogramming of the transcriptional landscape. Through a combination of long-term culturing, phenotypic and physiological assays, whole-genome sequencing, and transcriptomic analysis, we examined whether *S. eubayanus* undergoes a GASP-like evolutionary process and how these changes contribute to its cryotolerance. Our findings offer broader insights into the evolutionary significance of stationary-phase adaptation in wild yeasts and advance our understanding of microbial adaptation to extreme environments.

## Methodology

### Strain and Culture Conditions

Experiments were conducted using the haploid *S. eubayanus* strain H216.1 (*ho*Δ, *MAT*), which was previously generated from a wild-type diploid CL216.1 strain [39]. To initiate aging cultures, an overnight pre-inoculum was grown in YPD medium at 20 °C. From this saturated pre-inoculum, a 1:100 dilution was prepared in fresh YPD and divided into three independent culture tubes, each considered a separate “aging lineage.” The cultures were then maintained statically at 4 °C for up to 60 days without additional nutrient replenishment.

Sampling occurred at days 10, 20, 30, and 60. At each time point, the cell density in each lineage was estimated to determine appropriate plating dilutions. An aliquot from each aged culture was plated on YPD agar and incubated at 4°C for approximately 7 days to allow colony development. From each plate, the three largest colonies were selected for subsequent studies, yielding nine distinct aged isolates (three per lineage) for every sampling day. The selected colonies were grown at 20°C in YPD culture media and were stored in glycerol 20% at -80°C. A summary of the strains used in this work is described in **Table S1**.

### Growth Rate Assessment

To evaluate growth performance at low temperatures, parental and aged isolates were grown in YPD medium for 48 hours at 20°C to obtain pre-inocula. These cultures were then transferred into fresh YPD medium within 96-well microplates and incubated at 4 °C, with three technical replicates per isolate. Growth was monitored by measuring optical density at 600 nm (OD₆₀₀) at regular intervals using a Tecan microplate reader. The resulting growth data were analysed using the growthcurver R package [40], specifically employing the *SummarizeGrowthByPlate* function to calculate the area under the growth curve (AUC) for each replicate.

To quantify growth changes relative to the parental control, the AUC values for each aged isolate were normalized to the mean AUC of the non-aged parental strain, and the resulting values were expressed as log₂ fold-change (LFC). The statistical significance of differences between aged and control isolates was determined using two-sample t-tests, ANOVA, or Kruskal–Wallis tests, with appropriate post-hoc comparisons in R. Visualization and plotting of growth results were performed using the ggplot2 package [41].

### Freeze-Thaw Survival Assay

To assess freeze-thaw tolerance, parental and aged isolates were recovered from −80°C glycerol stocks and then spread onto YPD agar plates. A single colony from each isolate was used to grow a liquid YPD pre-inoculum. These pre-cultures were then used to inoculate fresh YPD medium and grown to the exponential phase (OD_600_ between 0.5 and 0.8).

Cell concentration was determined using a Neubauer hemocytometer, and cultures were diluted to obtain triplicate suspensions containing approximately 200 cells each. For each isolate, one set of triplicates was subjected to freezing at −20°C for 2 hours, while a parallel control set was held at 4°C [42]. Following the incubation, all samples were thawed at room temperature for 30 minutes, and the entire volume of each suspension was plated onto YPD agar. Colony Forming Units (CFUs) were counted after 24 hours of incubation. The percentage survival was calculated for each replicate using the formula:

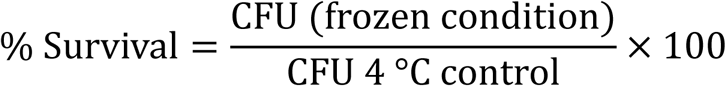

For comparative analysis, the survival percentage of each aged isolate was normalized to the mean survival of the non-aged parental strain. These ratios were then *log_2_*-transformed to calculate the LFC. A one-sample t-test was used to determine if the mean LFC of each aged cohort was significantly different from zero (i.e., different from the parental control). To assess significant differences among the aging time points, a Kruskal-Wallis test was performed, followed by a Dunn’s post-hoc test with Bonferroni correction for multiple comparisons. All statistical analyses and visualizations were performed in R.

### Fluctuation Assay

The mutation rate was determined using a fluctuation assay adapted from the protocol described by [43]. All steps of this assay were performed at 20 °C, the permissive, non-selective growth condition used throughout this study, to assess the inherent basal mutation rate of the parental and aged isolates. Initially, the three isolates, aged 20 and 60 days, along with the respective parental strain H216, were streaked onto YPD agar (2%) plates and incubated at 20 °C for approximately 48 hours. A single colony of each strain was then inoculated into ∼1 mL of SC-Arg medium and grown overnight at 20°C. Saturated cultures were diluted 1:10,000 in fresh SC-Arg medium, and 50 µL aliquots were distributed into 96-well microplates (n = 3 replicate plates per strain). The plates were sealed and incubated at 20 °C for 48 hours.

To select for mutants, 50 µL from each of the 72 wells were spotted onto SC-Arg-Ser medium with 60 mg/L canavanine and incubated at 20 °C for ∼120 h. To determine the total cell count (Nt), cultures from the remaining 24 wells were pooled, serially diluted, and plated in triplicate on YPD agar [44].

All data analysis was performed in R. The number of mutations per culture (m) was estimated from mutant colony counts using the *newton.LD* function from the *rSalvador* package [43, 45, 46]. The mutation rate was then calculated by dividing m by the total cell count (Nt) for each replicate. To quantify the effect of aging, the mutation rate of each aged isolate was normalized to the mean rate of the corresponding non-aged parental control (H216.1). These ratios were then log₂-transformed to calculate the LFC. The statistical significance of the LFC for each aged group (20 and 60 days) was determined using a one-sample t-test, comparing the mean LFC against a hypothetical mean of zero. Data was visualized using the ggplot2 package [41].

### Whole-Genome Sequencing and variant analysis

Genomic DNA of the 12 aged isolates (three per aged timepoint) was extracted from 5 mL overnight YPD cultures using the Zymo Quick-DNA bacterial/fungal extraction kit (cat. no. D6007, Zymo Research) following the manufacturer’s instructions. Library preparation and paired-end sequencing were performed on Illumina NextSeq500 in a Mid-Output Kit, with 150-bp paired-end reads, at the University of Santiago de Chile. Raw reads were inspected with FastQC v0.12.1 and adapter/quality trimmed with fastp v0.23.4 (-q 20 -f 15 -F 15). Trimmed reads were aligned against the haploid *S. eubayanus* CL216.1 [47, 48] reference genome with BWA-MEM algorithm using bwa software v0.7.18. Mapping statistics were obtained with Qualimap v2.2.2. SAM files were sorted, converted to BAM and tagged with read-group information using samtools v1.21 and samtools addreplacerg. PCR/optical duplicates were marked with GATK v4.6.1.0 (MarkDuplicates). All isolates were haploid; therefore, downstream ploidy was fixed to 1.

Variant calling followed GATK best practices adapted for haploids. First, per-sample gVCFs were generated with HaplotypeCaller (--sample-ploidy 1 --emit-ref-confidence GVCF --native-pair-hmm-threads 8). The raw VCFs were filtered with vcftools v0.1.16, retaining only SNPs that satisfied minQ ≥ 30, min-meanDP ≥ 10.

### RNAseq Analysis

Total RNA was extracted for transcriptomic analysis from *S. eubayanus* cultures representing two distinct conditions: the 60-day aged isolates and the parental (non-aged) control. For each condition, three biological replicates were grown in YPD medium at 4°C and harvested during the exponential growth phase. Prior to extraction, harvested cell pellets were washed twice with 50 mM EDTA and subjected to enzymatic digestion of the cell wall using Zymolyase (40 U) in Y1 buffer with beta-mercaptoethanol.

The RNA isolation was performed with the E.Z.N.A Total RNA Kit I (cat. no. R6834-02, Omega Bio-tek). The concentration and integrity of the resulting RNA were validated using a Qubit™ 4 Fluorometer (cat. no. Q33226; Thermo Fischer Scientific™) and an Agilent TapeStation system, respectively. For samples meeting the quality standards, cDNA libraries were prepared with the TruSeq Stranded Total RNA kit (cat. no. 20020598, Illumina). The prepared libraries were then sequenced on an Illumina NextSeq500 platform to yield 150-bp paired-end reads.

Raw FASTQ reads were quality-checked using FastQC (v0.12.1). Adapter sequences and low-quality bases were removed with Trimmomatic (v0.39), applying filtering parameters as follows: ILLUMINACLIP:TruSeq3-PE-2.fa:2:30:10 for adapter clipping, followed by SLIDINGWINDOW:4:20, LEADING:20, TRAILING:20, and MINLEN:36. The quality of the trimmed reads was subsequently re-assessed with FastQC. Trimmed, uncompressed reads were then aligned to the *S. eubayanus* CL216.1 reference genome using HISAT2 (v2.2.1). The genome index for HISAT2 was built incorporating splice site and exon information derived from the reference GTF annotation. Alignments were performed using paired-end mode, and the resulting SAM files were converted to sorted BAM format and indexed using Samtools (v1.19.1). Alignment summary statistics were generated by HISAT2 (**Table S2**).

Gene-level read counts were obtained from the sorted BAM files using the featureCounts function within the Rsubread R package (Bioconductor v2.16.1), based on the annotation file. Differential gene expression analysis between “Aged” and “Parental” conditions (n=3 replicates per condition) was conducted using DESeq2 (v1.42.0) in R. Genes with fewer than 10 total counts across all samples were excluded. Normalization and statistical testing were performed using default DESeq2 parameters, with “Parental” as the reference condition for comparison. Differentially expressed genes (DEGs) were identified based on an adjusted p-value (*p-adjusted*) < 0.05 and an absolute log2 fold change > 0.5. Volcano plots were generated using ggplot2 and ggrepel in R. Gene Ontology (GO) enrichment analysis for Biological Process (BP) was performed on up-regulated and down-regulated DEG lists against an appropriate background universe using clusterProfiler (v4.10.1) and the org.Sc.sgd.db annotation package (v3.18.0).

### Inference of Transcription Factor Activity and Regulatory Network Analysis

To identify potential transcriptional regulators (TFs) of DEGs, a transcription factor activity inference analysis was performed. Two primary evidence sources from the YeTFaSCo database were utilized: (i) expression enrichment data, which associates target gene expression with TF mutations, and (ii) TF binding motif enrichment data from the promoter regions of the genes of interest. For each TF, an integrated activity score was calculated by the evidence from expression data where the *p-values* were transformed using the formula −*log*_10_ (*p* +1e −300).

TFs were ranked based on their integrated activity score for both the up- and down-regulated gene sets. To visualize the regulatory network, TF-target gene interaction data were obtained from the YEASTRACT database. A network was constructed connecting the top 10 TFs with the highest activity score for the up-regulated genes to a curated set of target genes with high biological relevance to the stress-response phenotype. All analyses and visualizations were performed in R (v4.3.2) using the data.table package for robust data loading, tidyverse for data manipulation, and igraph and ggraph for network construction and visualization.

### Intracellular ROS Measurement

To quantify intracellular Reactive Oxygen Species (ROS), aged and parental isolates were first grown on YPD solid plates at 4°C. Biomass was then used to inoculate liquid YPD cultures, which were grown at 4°C with agitation. Cells were harvested from the cultures, centrifuged at 1700 xg for 4 min, and the resulting pellet was resuspended in 500 µL of phosphate buffer (80 mM Na_2_HPO_4_, 20 mM NaH_2_PO_4_, 100 mM NaCl, pH 7.4) containing 10 µM of the probe 2’,7’–dichlorofluorescin diacetate (DCFDA). Samples were incubated in darkness for 50 min at 20°C with agitation. After incubation, cells were pelleted again, washed, and resuspended in 500 µL of fresh phosphate buffer. 200 µL of each sample were transferred to a black 96-well optical-bottom plate, and fluorescence was measured from the bottom using a Cytation reader at an excitation/emission of 490/530 nm.

### Fluorescence Microscopy and Vacuolar Morphology

Yeast cells grown on solid YPD plates at 4°C were used to inoculate 0.5 mL of liquid YPD containing 3 µM of the vacuolar membrane dye FM™ 4-64 (Thermo Fisher). These cultures were incubated for 24 hours at 20°C with agitation and protected from light. For imaging, 0.2 µL of concentrated cells were mounted on a coverslip using immersion oil. Samples were observed on a Carl Zeiss 800 confocal microscope with a 63x objective at 2.5x zoom, using an excitation/emission of ∼565/744 nm. For each condition, 110-200 cells were imaged and manually classified into one of three phenotypes: Class A (single large vacuole), Class B (multiple small vacuoles), or Class C (highly fragmented) [49]

### Data Accessibility

All sequences have been deposited in the National Center for Biotechnology Information (NCBI) as a Sequence Read Archive under the BioProject accession number PRJNA1314239.

## Results

### Prolonged Aging at Low Temperature Confers Enhanced Fitness and Stress Resistance

To examine the phenotypic effects of long-term aging in *S. eubayanus*, aliquots from the 10-, 20-, 30-, and 60-day static cultures were plated. To identify the most fit individuals at each time point, isolates were sampled by deliberately selecting the three biggest colonies from each aged lineage (see Methods). These aged isolates were then phenotypically analyzed and compared to the non-aged parental strain. We will refer to ‘aged isolates’ as those recovered from aging cultures and subsequently regrown under non-selective, nutrient-rich conditions (see Methods). After recovering the aged isolates, we assessed the growth performance at the selective temperature of 4°C by quantifying the Area Under the growth Curve (AUC). While isolates from all aging time points displayed considerable phenotypic heterogeneity, the mean growth performance of isolates aged for 10, 20, and 30 days was not significantly different from that of the parental control (two-sample t-test; *p-value* > 0.05) (**Fig. 1A**). In contrast, isolates from the 60-day cohort exhibited a consistent and statistically significant increase in growth performance, with their AUC being, on average, 30% higher than that of the parental strain (mean LFC = 0.38; one-sample t-test, *p-value* <0.001) (**Fig. 1A**). A one-way ANOVA further confirmed a significant effect of aging time on growth performance (*p-value* <0.001), and post-hoc analysis revealed that the growth of the 60-day cohort was significantly higher than that of all other time points (*p-value* <0.001).

**Fig. 1.**
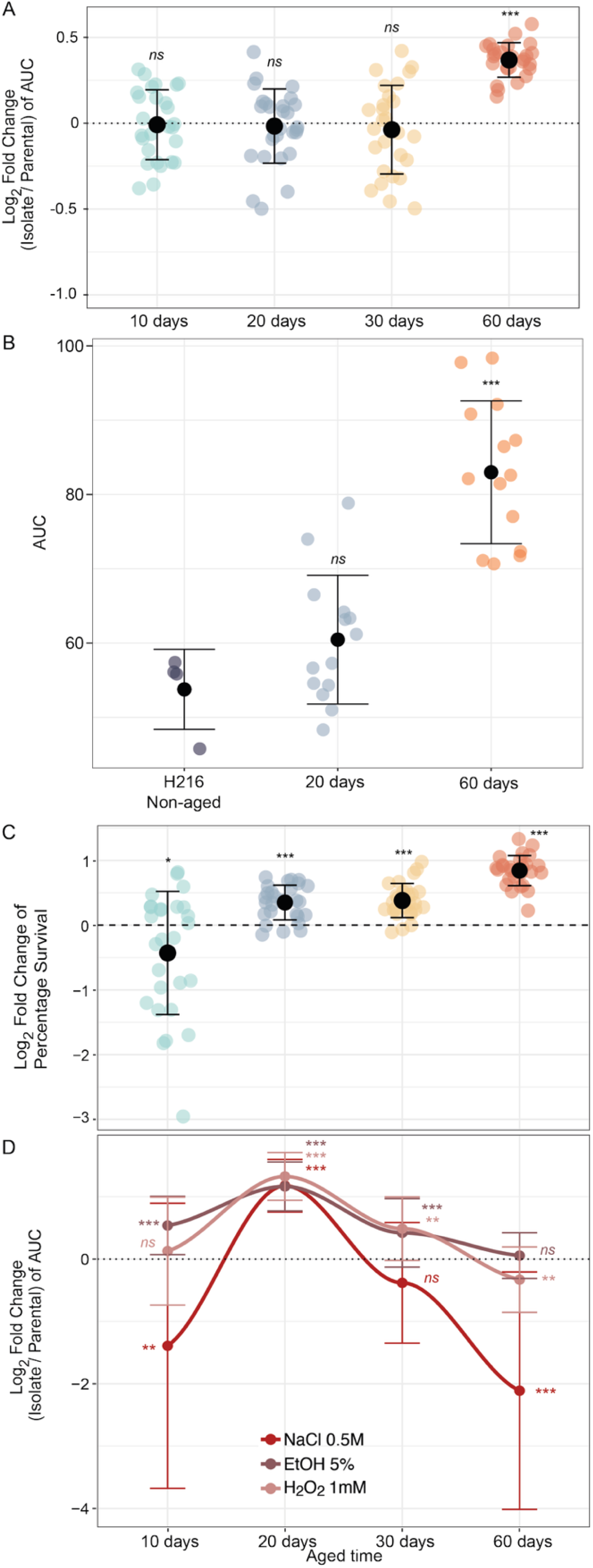
Growth capacity of *S. eubayanus* isolates during long-term aging at 4°C. (A) Growth performance at 4°C. Data points represent the log₂ fold change (LFC) of the area under the curve (AUC) for individual aged isolates relative to the mean AUC of the non-aged parental control (dashed line at LFC = 0). Black dots and error bars show the mean ± standard deviation for each aging cohort (10, 20, 30, and 60 days). (B) Stability of the enhanced growth phenotype. Growth performance (AUC) at 4°C was re-assessed after parental, 20-day, and 60-day isolates were serially passaged for ∼40 generations under non-selective conditions (YPD at 20°C). Data points show raw AUC values, confirming the stable, transgenerational nature of the adaptation in the 60-day cohort. (C) Freeze-thaw tolerance. Data points represent the LFC of survival percentage for individual-aged isolates relative to the parental control. (D) Cross-protection against other abiotic stressors. The LFC of AUC is shown for isolates from different aging time points when grown in the presence of 0.5 M NaCl, 5% ethanol (EtOH), or 1 mM H₂O₂. Points and error bars represent the mean ± standard deviation of biological replicates for each cohort. Asterisks (*) above the data points indicate statistically distinct groups as determined by one-way ANOVA followed by Tukey’s HSD post-hoc test (* *p-value* < 0.05, ** *p-value* < 0.01 and *** *p-value* < 0.005).

To determine whether this enhanced fitness was a stable transgenerational trait rather than a transient short physiological memory, isolates from the 20- and 60-day cohorts were serially passaged for seven passages (approximately 40 generations) under non-selective conditions at 20°C. Following this period, their growth performance was re-assessed at the selective temperature of 4°C. Notably, the 60-day isolates fully retained their significant growth advantage over the parental strain, suggesting that the acquired phenotype is a stable trait (**Fig. 1B**). In contrast, the 20-day isolates showed no significant difference from the parental control after the passages, consistent with the original observation.

Next, we evaluated the tolerance of the isolates to physical stress using a freeze-thaw survival assay, which revealed a complex, time-dependent response. Initially, the 10-day aged cohort exhibited a significant reduction in survival compared to the parental control (*p-value* = 0.03; **Fig. 1C**). In contrast, all subsequent cohorts demonstrated enhanced fitness, with isolates from 20, 30, and 60 days showing a marked and significant improvement in freeze-thaw tolerance (*p-value* < 0.001 for all; **Fig. 1C**). Inter-group comparisons further delineated this progressive adaptation: while the 20- and 30-day cohorts performed similarly, both exceeding the 10-day cohort. The 60-day cohort ultimately achieved the highest level of tolerance, significantly outperforming all other groups (ANOVA with Tukey’s HSD, *p-value* < 0.05). These results indicate a greater freeze-thaw tolerance in 60-day aged isolates relative to any other cohort.

To determine whether the acquired cold tolerance provided a broader cross-protection phenotype or a fitness trade-off, we evaluated the growth (measured as AUC) of aged isolates under abiotic stressors: 0.5 M NaCl, 5% EtOH, and 1 mM H₂O₂. A two-way ANOVA revealed significant main effects for both aging time (F(4, 388) = 48.78, *p-value* < 0.001) and stressor type (F(2, 388) = 53.22, *p-value* < 0.001). We found a highly significant interaction between these two factors (aging time × stressor, F(8, 388) = 6.71, *p-value* < 0.001), indicating that the adaptive response to aging is stressor-dependent (**Fig. 1D**). For example, post-hoc analysis showed that isolates aged for 20 days consistently displayed the highest level of cross-protection. This group exhibited significantly improved growth tolerance compared to the non-aged control across all three tested conditions (one-sample t-tests with BH correction, *p-adjusted* < 0.001 for NaCl, EtOH, and H₂O₂), exhibiting a more generalist-like behaviour. In contrast, the 60-day aged isolates did not exhibit a general tolerance. In fact, their growth was significantly impaired under saline and H₂O₂ stress compared to the control (*p-adjusted* < 0.001 for NaCl and *p-adjusted* = 0.006 for H₂O₂; **Fig. 1D**), exhibiting a more specialist-like behaviour. The loss of tolerance was particularly evident for NaCl, where the 60-day isolates performed significantly worse than all other aging periods, including the 30-day isolates (Tukey’s HSD, *p-adjusted* < 0.001). Furthermore, at the 60-day time point, tolerance to NaCl was significantly lower than to either EtOH or H₂O₂ (*p-adjusted* < 0.001 for both comparisons).

Taken together, these results demonstrate that prolonged aging selects for fitter isolates with enhanced specific cold resistance, suggesting trade-offs between traits. Furthermore, they reveal that the adaptive trajectory towards this specialization is complex and time-dependent, differing from the path for general stress cross-protection, which instead peaks at an intermediate stage of 20 days.

### Aged Isolates Exhibit Altered Redox State, Vacuolar Morphology, and a greater mutation rate

Based on the distinct growth and stress tolerance phenotypes observed in the initial screening, we selected the 20-day and 60-day aged cohorts for further physiological characterization to assess their cellular state after prolonged cold exposure. We investigated their intracellular redox environment and vacuolar integrity, two key indicators of cellular homeostasis. To evaluate the oxidative state, we quantified intracellular ROS levels. Both the 20-day and 60-day aged populations displayed a statistically significant increase in mean ROS levels compared to the non-aged parental control (mean LFC = 0.43, one-sample t-test, *p-value* < 0.001, mean LFC = 0.47, one-sample t-test, *p-value* < 0.001, respectively) (**Fig. 2A**). While there was considerable variation among individual isolates within the aged cohorts, the overall trend indicated a constitutively higher oxidative state in cells that had undergone long-term aging.

**Fig. 2.**
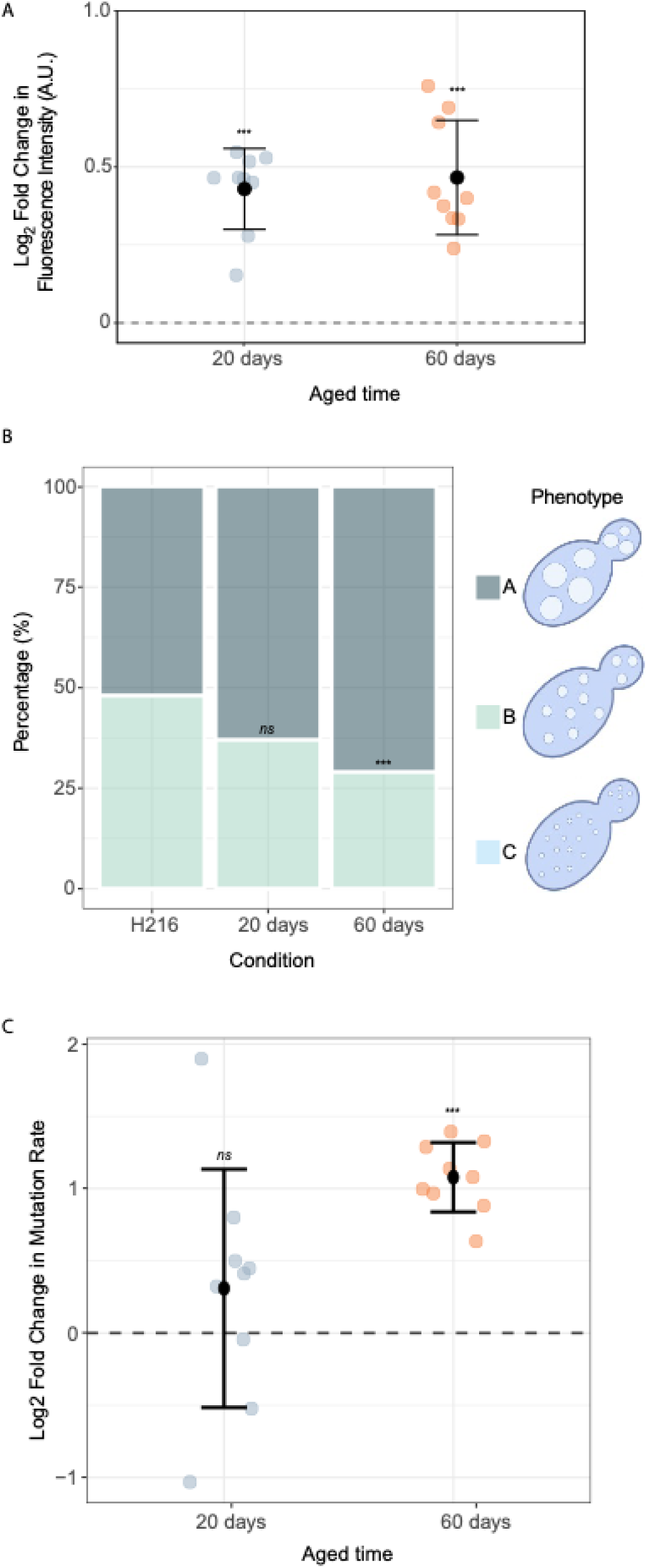
Characterization of aged *S. eubayanus* isolates. (A) Intracellular ROS levels. Data points represent the LFC in relative fluorescence units for individual isolates from the 20-day and 60-day aged cohorts, relative to the mean of the parental control (H216, represented by the dashed line at LFC=0). Black points and error bars indicate the mean ± standard deviation for each cohort. (B) Distribution of vacuole morphologies. Stacked bar chart shows the mean percentage of cells from each cohort exhibiting Class A (large, single vacuole), Class B (multiple small vacuoles), or Class C (highly fragmented) phenotypes, as determined by fluorescence microscopy. Representative schematics for each vacuole class are shown on the right. Statistical significance of the distribution shift compared to the H216 control was determined by a Chi-squared test. (C) LFC in the mutation rate of isolates aged for 20 and 60 days, relative to the non-aged parental control (dashed line at LFC=0). Each point represents an independent aged isolate. Black dots and error bars indicate the mean ± standard deviation. *P-values* were calculated using a one-sample t-test against a hypothetical mean of zero.

Next, given that vacuolar morphology is a critical indicator of cellular health and stress response [50, 51], we analysed the distribution of vacuolar phenotypes using FM4-64 staining. We classified the cells into three categories, labelled A to C, with A representing the healthiest vacuolar morphology. The parental strain exhibited a baseline distribution, with approximately half of the cells (48%) displaying multiple small vacuoles with defined borders (Class B). In contrast, the aged cohorts showed a clear shift towards the non-fragmented Class A morphology (**Fig. 2B**). While the increase in the proportion of Class A cells in the 20-day cohort was not statistically significant compared to the parental control (Chi-squared test, *p-value* = 0.067), the shift was highly significant for the 60-day cohort (Chi-squared test, *p-value* < 0.001). This latter group was predominantly composed of cells with the Class A vacuolar phenotype, indicating a shift towards a cellular organization characteristic of entry into quiescence [18, 19]compared to non-aged cells.

To assess whether aged cells exhibited a greater mutation rate than non-aged cells, we determined the mutation rates of 20-day and 60-day cohorts, as well as those of non-aged parents, using a canavanine fluctuation assay. 20-day-old isolates exhibited no significant change in their mutation rate compared to the non-aged parental control (mean LFC = 0.31; one-sample t-test, *p-value* = 0.29; **Fig. 2C**). In contrast, a significant increase in the mutation rate was observed in the 60-day-old isolates. Isolates from this group showed an average mutation rate ∼ 6.89 x 10^-7^ mutations per division) approximately double that of the parental control (∼ 3.22 x 10^-7^ mutation per division), representing a highly significant change (mean LFC = 1.08;one-sample t-test, *p-value* < 0.001; **Fig. 2C**). These findings would be consistent with a GASP phenotype, where aging in culture media without nutrient replenishment progressively could compromise genomic integrity.

### Genomic Variant Dynamics Reveal a Pattern of Fluctuation and Selection During Long-Term Aging

To identify potential genetic changes in aged cells underlying the cold-growth improvement, we sequenced the genomes of twelve selected isolates, three from each aging time point (10, 20, 30, and 60 days). Contrary to a simple model of steady mutation accumulation, our analysis revealed a dynamic pattern of genomic variation characterized by an initial burst of mutations followed by selection. The highest number of single-nucleotide polymorphisms (SNPs) was detected in the 10-day cohort, after which the variant count decreased sharply (**Fig. 3A, Table S3**). By day 60, the three sequenced isolates collectively harboured only five SNPs. Despite the elevated mutation rate (Fig. 2C), most variants did not persist, suggesting that only a few genotypes were maintained. These variants were distributed across the genome, with some regions, such as on chromosome II, appearing as potential mutational hotspots, where multiple variants arose in independent aging lineages and time points (**Fig. 3A**, **Table S3**).

**Fig. 3.**
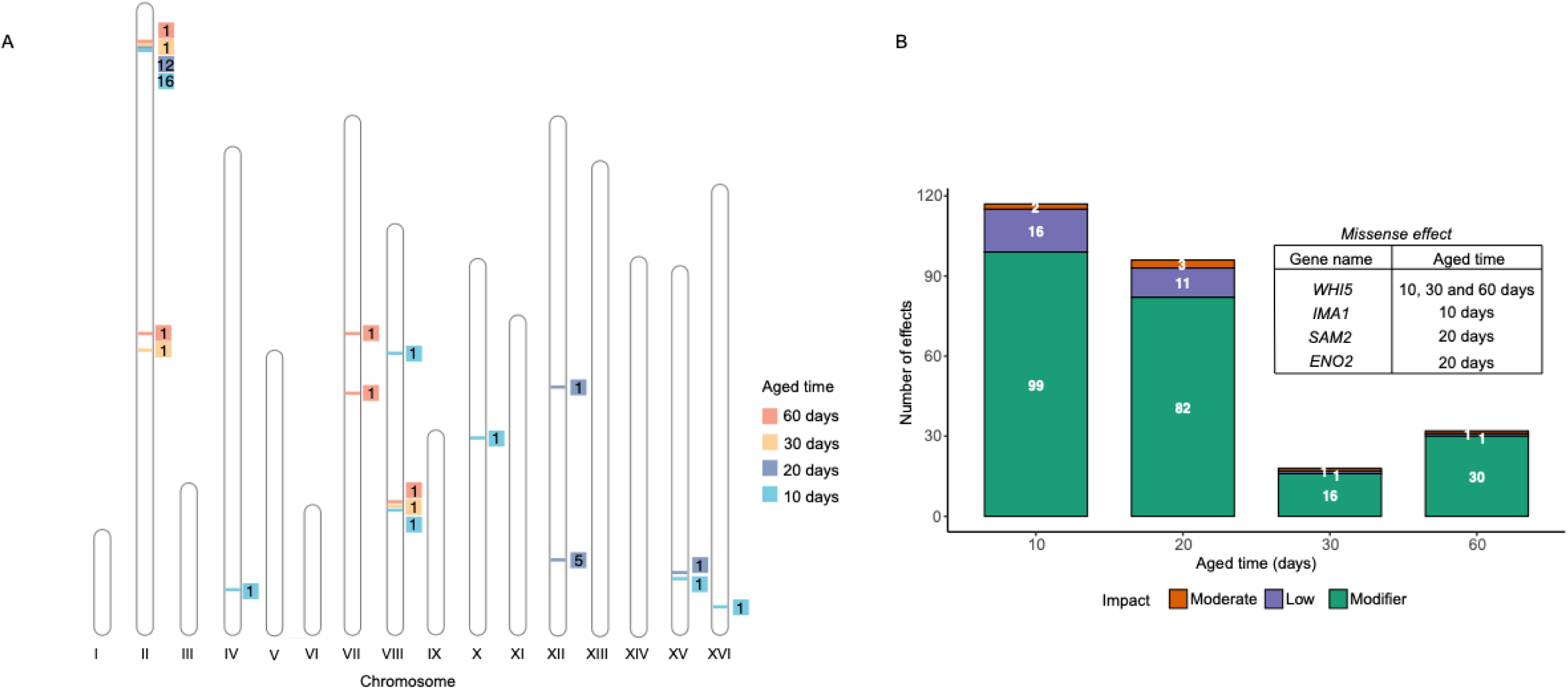
Genomic and changes in aged cells. **(A)** Chromosomal distribution of identified SNPs across the four aging time points (10, 20, 30, and 60 days). Colours indicate the time point at which a SNP was first detected. Numbers adjacent to highlighted chromosomal bands denote the total count of unique SNP locations identified within that specific region. **(B)** Stacked bar chart quantifying the total number of variants per aging cohort, categorized by their SnpEff-predicted impact (Modifier, Low, Moderate). The inset table lists recurrent missense mutations identified in key genes.

We classified the potential functional consequences of these variants by using SnpEff. The total number of variants, particularly those categorized as having ‘Modifier’ (e.g., variants in non-coding regions) or ‘Low’ impact (e.g., synonymous changes), peaked at day 10 and subsequently decreased sharply (**Fig. 3B**). Particularly noteworthy was the identification of a recurrent missense mutation in *WHI5*, a master repressor of the G1/S cell cycle transition. Mutations in *WHI5* were independently detected in isolates from 10, 30, and 60 days in the same-aged line, and their presence was confirmed by Sanger sequencing. While this pattern suggests a strong selective pressure, it was only detected in one of the three independent lineages sequenced at day 60 (**Table S3**). However, because all isolates exhibited enhanced fitness (Fig. 1A), this single mutation is unlikely to account for the environmental adaptation, suggesting alternative mechanisms. Other mutations in metabolic and regulatory genes, such as *IMA1*, *ENO2*, and *SAM2,* appeared only transiently at earlier time points and were not detected in the final adapted populations (**Fig. 3B**, inset table). This pattern of mutation fluctuation, combined with the absence of recurrent variants across lineages, strongly indicates that the accumulation of point mutations does not mainly shape the long-term adapted state.

### Transcriptional reprogramming underlies the improvement in fitness of aged cells

We hypothesize that a more immediate and plastic mechanism likely drives the robust adaptive changes in aged cells. Therefore, we next investigated the molecular basis of the adapted phenotype by analysing the global transcriptional reprogramming in the 60-day isolates. For this, we compared the transcriptome of 60-day aged isolate (**Table S2**) to the parental control during exponential growth at 4°C. Principal Component Analysis (PCA) of the transcriptomes revealed a distinct separation between the aged and parental samples, with the first principal component (PC1) accounting for 80% of the dataset’s variance (**Fig. S1**). The analysis identified 256 differentially expressed genes (DEGs) (*p-adjusted* < 0.05, |log₂FC| > 0.5), with 164 genes up-regulated and 92 down-regulated in the aged cells relative to the parental control (**Fig. 4A, Table S4**).

**Fig. 4.**
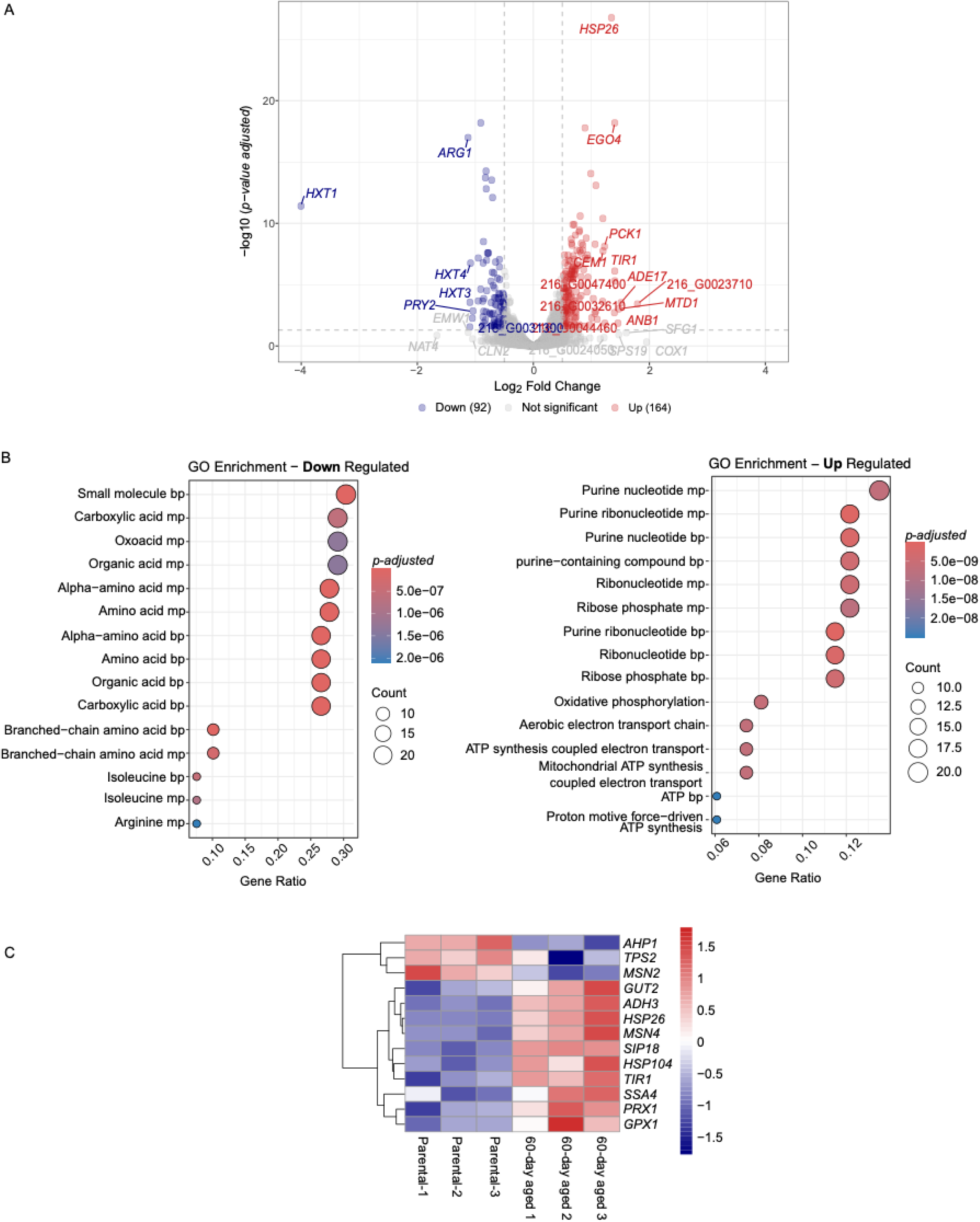
Genomic and Transcriptional changes in aged cells. **(A)** Volcano plot illustrating differentially expressed genes. Points are coloured based on statistical significance (*p-adjusted* < 0.05) and LFC (>|0.5|). **(B)** Gene Ontology (GO) enrichment dot plots for Biological Process (BP) terms of down-regulated (left) and up-regulated (right) genes in 60-day aged isolates versus the parental control. Dot size corresponds to the number of genes (Count) in each term, and colour represents the adjusted *p-value*. **(C)** Heatmap displaying the LFC of differentially expressed genes (*p-adjusted* < 0.05) from a curated list of cold and stress response-related genes (**Table S5**). Columns represent biological replicates, and rows represent individual genes.

Gene Ontology (GO) enrichment analysis provided insight into the processes altered in the adapted cells. Up-regulated genes were significantly enriched for Biological Process (BP) terms related to energy production and precursor synthesis, including “oxidative phosphorylation” (*p-adjusted* < 0.001), “ATP biosynthetic process” (*p-adjusted* < 0.001), and “purine ribonucleotide biosynthetic process” (*p-adjusted* < 0.001) (**Fig. 4B**). Consistent with these findings, Cellular Component (CC) analysis showed a strong enrichment for mitochondrial localizations, such as “mitochondrial inner membrane” and “respirasome” (**Fig. S2A**). The corresponding Molecular Function (MF) analysis highlighted terms like “transmembrane transporter activity”, “cytochrome−c oxidase activity” and “oxidoreductase activity” (**Fig. S2B**). Conversely, down-regulated genes were primarily enriched for BP terms associated with broad anabolic pathways, including “small molecule biosynthetic process” and, more specifically, “amino acid biosynthetic process” (**Fig. 4B**). The MF analysis of this set revealed enrichment for terms such as “ligase activity” and various carbohydrate “transporter activities” (**Fig. S2C**). In contrast, no significant enrichment for CC terms was detected.

To specifically probe the expression of genes with known roles in cold adaptation and related stresses, we examined a curated list of candidates known to have a role in cold tolerance (**Table S5**). A heatmap of the DEGs from this list revealed a distinct and consistent expression signature separating the aged and parental replicates (**Fig. 4C**). Notably, several key stress-responsive genes, including the heat shock proteins *HSP26* and *HSP104*, the master stress transcription factors *MSN2* and *MSN4*, and antioxidant enzymes such as *AHP1*, *PRX1,* and *GPX1*, were among those significantly up-regulated in the 60-day aged isolates. This pattern indicates a clear transcriptional reprogramming for cold tolerance in aged cells relative to non-aged cells.

To identify the transcriptional regulators responsible for this adaptive expression profile, we inferred TF activity by combining evidence from expression and motif enrichment data. This analysis revealed a distinct set of master regulators likely driving the up-regulated gene response. The TFs Usv1, Hac1, Sko1, Hot1, Rsc3, Gcr1, Msn4, Msn2, Cad1, and Pho4 emerged as the top ten candidates with the highest activity scores (**Fig. 5A**).

**Fig. 5.**
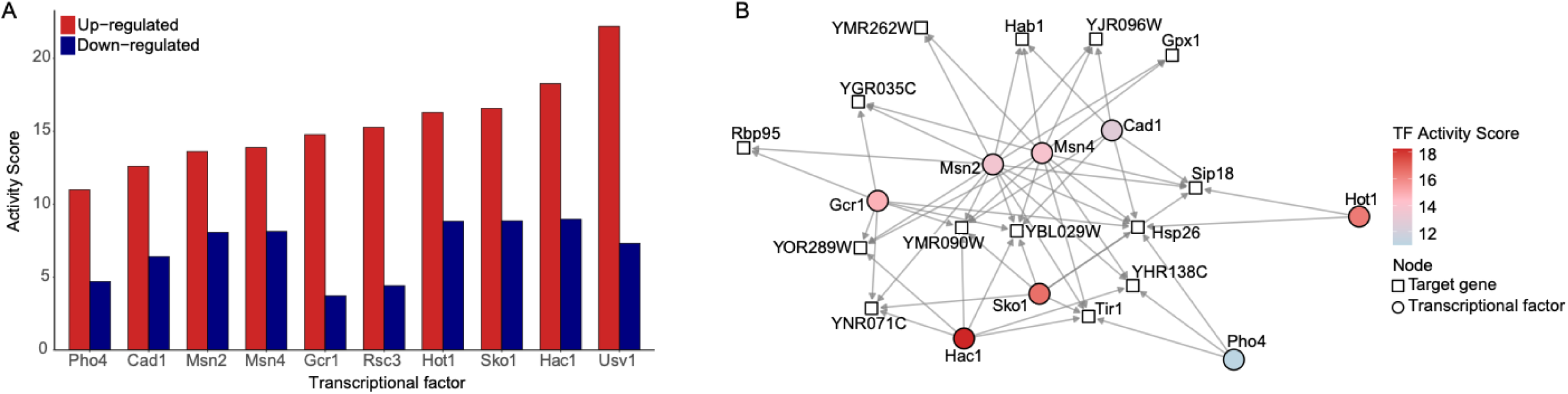
Inference of the Transcriptional Regulatory Network Driving the Adapted Phenotype. (A) Bar chart showing the top ten transcription factors (TFs) ranked by their activity score for the up-regulated gene set (red bars). The corresponding scores for the down-regulated gene set are shown for comparison (blue bars). Scores were calculated by integrating evidence from expression and motif enrichment data. (B) Gene regulatory network connecting the top-ranked TFs (circles) to a curated set of key stress-responsive target genes (squares). The color of each TF node corresponds to its activity score, as indicated in the legend. Edges represent documented regulatory interactions from the YEASTRACT database.

To visualize the regulatory connectivity of these top-ranked TFs, we constructed a gene regulatory network (GRN) linking them to a curated set of key stress-responsive target genes previously shown to be up-regulated (**Fig. 4C**). The resulting network reveals a highly interconnected regulatory hub where central stress effectors converge to control critical cellular defence genes (**Fig. 5B**). Notably, the paralogs Msn2 and Msn4, the canonical mediators of the General Stress Response, alongside the HOG pathway regulators Sko1 and Hot1, were identified as key nodes regulating the expression of genes encoding chaperones (*HSP26*, *HSP104*) and antioxidant enzymes (*GPX1*, *PRX1*). This visualization provides a qualitative map of the core pro-survival circuitry activated in the 60-day adapted isolates.

## Discussion

### Long-Term Cold survival Drives Adaptive Evolution in *S. eubayanus*

Our findings demonstrate that prolonged incubation under cold, nutrient-limiting conditions provides a significant adaptive response in *S. eubayanus*. Here, we found that long-term cold adaptation is driven by transcriptional reprogramming, mostly linked to oxidative stress resistance. This process, observed within a CLS experimental framework, partly shares key features with the GASP phenomenon, a well-documented survival strategy in bacteria like *E. coli* [5–8]. The GASP phenotype is defined by the emergence of mutants that can outcompete the ancestral population by more efficiently scavenging nutrients released from dead cells [6, 52, 53]. Indeed, functionally similar processes, often termed “adaptive regrowth,” have been described in yeast CLS experiments, where fitter mutants can emerge and take over the culture [38, 54, 55].

The isolates from day 60 clearly exhibited enhanced fitness, showing superior growth at the selective temperature of 4°C. Our whole-genome sequencing revealed a low number of SNPs. This modest genomic footprint suggests that the accumulation of numerous point mutations does not necessarily drive fitness gains. We demonstrate that the enhanced growth phenotype is a stable, transgenerational trait, which is fully retained after seven consecutive passages under non-selective conditions. This finding allows us to rule out transient physiological memory and suggests that the stable transcriptional reprogramming itself is the basis for the adaptation. This leads us to speculate that the primary engine of adaptation in this system is the extensive and heritable transcriptional reprogramming observed in the aged isolates. This large-scale rewiring of gene expression, characterized by a shift towards energy production and a robust stress response, enables a more efficient physiological state. This constitutes a different adaptive path than a classic, mutation-driven GASP takeover. Such a strategy relying on stable transcriptional states is ecologically significant for *S. eubayanus*, a species native to the fluctuating environments of Patagonia [25, 29].

### The Dynamics of Adaptation Reveal a Fitness Trade-Off Between Generalism and Specialism

A temporal analysis of the evolving populations reveals a classic evolutionary fitness trade-off, where adaptation to one environment occurs at the expense of fitness in others [56, 57]. Such trade-offs are a common, though not universal, outcome in microbial evolution experiments [56, 58]. In our study, while fitness in the selective cold environment progressively increased, tolerance to other stressors followed a non-linear path. At the 20-day intermediate stage, the isolates displayed a “generalist” phenotype, characterized by enhanced cross-protection against high salinity, ethanol, and oxidative stress. This suggests an initial adaptation based on broad, likely costly, stress defense mechanisms [59–62]. However, by day 60, this generalism was lost. The highly cold-adapted isolates became specialists, showing significantly decreased tolerance to NaCl and H_2_O_2_ stress compared to the ancestor.

This transition from a generalist to a specialist strategy is a microevolutionary example of a dominant macroevolutionary pattern observed across the Saccharomycotina subphylum [63, 64]. Genome-scale analyses have shown that yeast diversification is primarily characterized by reductive evolution, where lineages frequently narrow their metabolic niche breadth and become more specialized [64, 65]. As selection continued under constant cold and starvation in our experiment, the populations shed what became metabolically expensive, broad-spectrum defences in favour of a more efficient physiology optimized for the specific conditions. It is noteworthy that this specialization, resulting from the loss of plasticity, does not imply that generalists are inherently less efficient; in fact, across yeasts, metabolic breadth often correlates positively with growth efficiency and a more versatile genetic toolkit [66]. This trajectory underscores the principle that adaptation is usually constrained by antagonistic pleiotropy and physiological trade-offs [67, 68].

### Physiological Reprogramming Supports the Adapted State

The observed adaptive phenotypes are supported by a fundamental, likely epigenetic reprogramming of cellular physiology that is heritable. A key feature of the aged cells is a significant increase in intracellular ROS. While high ROS levels are widely associated with the damaging effects of aging due to the accumulation of oxidative damage to macromolecules [69, 70], a more nuanced view frames ROS as dual-function molecules [71, 72]. At moderate levels, ROS can act as signalling molecules that trigger protective pathways, a phenomenon known as mitohormesis [69, 71, 73]. The elevated ROS in our 60-day isolates, paired with their enhanced fitness, supports this dual role. The sustained oxidative environment likely acts as both a mutagenic force, contributing to the increased mutation rate by causing DNA damage [74], and as a constant signal that maintains a robust, constitutive stress defence, evidenced by the upregulation of antioxidant genes.

A second critical adaptation is the dramatic reorganisation of the vacuole. The transition from fragmented vacuoles to a single, large “Class A” morphology in the 60-day isolates is a well-established cytological marker of entry into a quiescent, long-lived state [50, 75]. This morphology is optimal for large-scale degradation and nutrient recycling via autophagy, which is essential for survival during starvation [54, 76]. This structural change aligns perfectly with the physiological requirements of a cell adapted for long-term survival and scavenging.

### Molecular Basis of a Cold-Adapted, Scavenging Lifestyle

The genetic basis of the adapted phenotype appears complex and not driven by the accumulation of numerous point mutations. Despite the low overall number of SNPs detected, the identification of a recurrent missense mutation in *WHI5* is particularly noteworthy (**Table S4**). As the yeast functional analogue of the human Retinoblastoma (Rb) protein, Whi5 is a master repressor of the G1/S cell cycle transition and a key gatekeeper of quiescence [77–83]. An adaptive mutation in such a crucial regulator is therefore a plausible contributor to the observed pro-survival phenotype. However, it is unlikely that this single genetic change can fully account for the profound and system-wide adaptive changes observed. Instead, the most comprehensible molecular blueprint for the adapted state was revealed at the transcriptomic level. The global expression profile of our 60-day isolates is the canonical hallmark of entry into quiescence, representing a strategic turn away from proliferation and towards long-term survival, a state strongly associated with extended CLS [18, 37, 84, 85]. This is evidenced by two key observations. First, we see a significant repression of anabolic processes, particularly amino acid biosynthesis, a classic outcome of TORC1 inhibition that signals a strategic change from proliferation to resource conservation [13, 86]. Second, this is accompanied by a striking upregulation of genes whose functions are predominantly localized to the mitochondrion, with GO terms for “oxidative phosphorylation” and “ATP biosynthetic process” being highly enriched. This metabolic shift towards highly efficient aerobic respiration is representative of longevity in yeast, allowing cells to efficiently scavenge and utilize the scarce resources released by dead cells [12, 87]. The inferred attenuation of TORC1/PKA signalling is known to activate the General Stress Response (GSR) by promoting the nuclear localization of the transcription factors *MSN2* and *MSN4*. Consistent with this, we observe a robust upregulation of canonical *MSN2*/*4* target genes, including molecular chaperones (Hsp26, Hsp104) and antioxidant enzymes (Prx1 and Gpx1), which are essential for coping with the increased oxidative stress inherent to both aging and enhanced respiration [36, 88, 89]. Our transcription factor activity analysis provides further mechanistic insight into reprogramming. The inference confirmed the central role of the General Stress Response, identifying Msn2 and Msn4 as key TFs with the highest activity scores for the upregulated gene set and central to the GRN. Both TFs are highly associated with transcriptional response under various stresses [36, 60, 90]. Additionally, key regulators of the HOG pathway, Sko1 and Hot1, were also identified. Interestingly, Usv1, a TF known in *S. cerevisiae* for its role in osmotic stress response and growth on non-fermentable carbon sources [91, 92], also ranked highly. However, its connections to our curated network of stress genes were not apparent, perhaps pointing to a distinct or expanded regulatory role in *S. eubayanus*. Collectively, the coordinated activity of this suite of master regulators provides the mechanistic basis for the shift towards a robustly protected, quiescent state. Thus, this system-wide transcriptional reprogramming, resulting in a stable and heritable pro-survival state maintained by attenuated pro-growth signalling, provides the robust molecular foundation for the superior fitness and cryotolerance of the 60-day aged isolates.

## Conclusion

In this study, we demonstrate the mechanism by which a cryotolerant wild yeast overcomes to prolonged cold and starvation. Our integrated analysis reveals a multifaceted adaptive strategy, with the most profound changes observed at the transcriptional level. We show that a massive, transgenerational reprogramming shifts the cellular economy from anabolism towards a catabolic, scavenging state powered by enhanced mitochondrial respiration. This stable transcriptional state is the primary engine of adaptation. Concurrently, we identified a modest number of genomic changes, including a recurrent and likely adaptive mutation in the cell cycle regulator *WHI5*. Although the exact relationship between the limited genetic mutations observed and the overall change in gene expression is not yet fully understood, our results clearly demonstrate that long-term survival and increased fitness during extended cold and nutrient scarcity are achieved through a stable, reprogrammed transcriptional state. This work highlights that non-mutational mechanisms can serve as a powerful and primary driver for microbial adaptation to persistent environmental stress.

## Author contribution

Conceptualization, L.A.S. and F.A.C.; methodology, L.A.S., M.LH., T.M.C., P.V., V.Z. and F.A.C.; formal analysis, L.A.S. and F.A.C.; investigation, L.A.S.; writing—original draft preparation, L.A.S. and F.A.C.; writing—review and editing, L.A.S., P.V. and F.A.C.; funding acquisition, L.A.S. and F.A.C. All authors have read and agreed to the published version of the manuscript.

## Declaration of competing interest

The authors declare that they have no competing interests.

## Acknowledgements

L.A.S. acknowledges ANID POSTDOCTORAL FONDECYT [3240351]. F.A.C acknowledges ANID FONDECYT [1220026] and ANID - Programa Iniciativa Científica Milenio - ICN17_022 and NCN2024_040. PV acknowledges FONDECYT INICIACIÓN [11240649], and ANID - Programa Iniciativa Científica Milenio ICN17_022 and NCN2021_050. MLH acknowledges Universidad de Santiago de Chile POSTDOC_DICYT, Código 022443CR_Postdoc Vicerrectoría de Investigación, Innovación y Creación. We thank Gilles Fischer for his constructive feedback on the manuscript.

## Data availability

All raw data from the metabarcoding sequencing have been deposited in the Sequence Read Archive (SRA) of the National Center for Biotechnology Information (NCBI) under BioProject accession number PRJNA1320675

